# Hyperspectral imaging for chloroplast movement detection

**DOI:** 10.1101/2024.03.26.586650

**Authors:** Paweł Hermanowicz, Justyna Łabuz

## Abstract

We employed hyperspectral imaging to detect chloroplast positioning in *Nicotiana benthamiana* and *Arabidopsis thaliana* leaves and assess its influence on commonly used vegetation indices. In low light, chloroplasts move to cell walls perpendicular to the direction of the incident light. In high light, they move to cell walls parallel to the light direction. Chloroplast movements result in significant changes in leaf transmittance and reflectance. The changes in leaf reflectance offer a way to examine chloroplast positioning in a non-contact way. At the same time, they may confound remote sensing of other physiological traits. The shape of reflectance spectra recorded on irradiated and non-irradiated parts of *N. benthamiana* and *A. thaliana* leaves indicated the specific position of chloroplasts. Low blue light resulted in a decrease in leaf reflectance in the green-yellow region of the spectrum. High blue light irradiation caused an increase in leaf reflectance in the visible range. The differential spectra, showing the effect of high light on leaf reflectance, exhibited a characteristic saddle in the green-yellow region and a peak at around 695 nm. Results obtained for A. thaliana mutants with disrupted chloroplast movements suggest that the observed spectral changes are mostly due to the chloroplast relocations. The reflectance spectra were used to train machine learning methods in the classification of leaves according to the chloroplast positioning. The convolutional network showed low levels of misclassification of leaves irradiated with high light even when different species were used for training and testing. This suggests that reflectance spectra may be used to detect the chloroplast avoidance response in heterogeneous patches of vegetation. We also examined the correlation between chloroplast positioning and values of indices of normalized-difference type for various combinations of wavelengths and proposed a chloroplast movement index for validation of chloroplast positions in leaves. The analysis of commonly used vegetation indices showed that their values may be altered due to chloroplast rearrangements. Our work indicates that changes in leaf reflectance due to chloroplast movements may be substantial and should be taken into account in remote sensing studies.

## Introduction

Plants show a remarkable ability of chloroplast relocation in changing light conditions, for review see (Łabuz et al., 2022). In low light, chloroplasts move to cell walls perpendicular to the direction of incident light. This response, called chloroplast accumulation, increases light capture for photosynthesis (Gotoh et al., 2018). In high light, chloroplasts gather at cell walls parallel to the direction of incident light. Chloroplast avoidance protects the photosynthetic apparatus from excess light (Kasahara et al., 2002; Sztatelman et al., 2010), (Davis & Hangarter, 2012) or shapes light gradients within the leaf tissue (Brugnoli & Bjorkman, 1992), (Wilson & Ruban, 2020). Seed plants move chloroplasts in response to blue light, however, red light is also active in some ferns, mosses, and green algae (Suetsugu, et al., 2005). In the model plant *Arabidopsis thaliana*, blue light-induced chloroplast movements are controlled by photoreceptor serine-threonine kinases called phototropins (Christie, 2007). Two phototropins, phot1 and phot2, mediate the chloroplast accumulation response (Sakai et al., 2001). Sustained chloroplast avoidance can be elicited only by phot2 (Jarillo et al., 2001), (Kagawa et al., 2001). Thus, no substantial directional chloroplast movements are observed in the *phot1phot2* double phototropin mutant. The *phot1* mutant is less sensitive to very low light irradiances than wild-type plants, but it shows both accumulation and avoidance response (Luesse et al., 2010). In the *phot2* mutant, light elicits chloroplast accumulation regardless of irradiance, though it is preceded by a weak, transient avoidance response in high light (Luesse et al., 2010), (Łabuz et al., 2015). Chloroplast avoidance is observed in both low and high light in *jac1,* a signaling mutant, lacking JAC1, the J-domain protein required for chloroplast accumulation response 1 (Suetsugu, et al., 2005).

First microscopic chloroplast movement observations were performed by Senn (Senn, 2008), revealing a plethora of different arrangements within plant cells (Kataoka, 2015). Microscopic observations still play a pivotal role in assessing chloroplast positions within leaf cells, but they are laborious (Kagawa & Wada, 2000) and provide information for individual cells. Interpretation of microscopic observations requires care due to the steep fluence rate gradient within the mesophyll tissue. In some species, the effects of chloroplast movements on the optical properties of the leaves are so pronounced that they may be observed with the naked eye. Whereas chloroplast avoidance results in paler leaf blades, chloroplast accumulation causes the darkening of leaves. This effect is the basis for the so-called slit assay (Kagawa & Wada, 2000), and is even utilized by artists (Dolinsky & Hangarter, 2013). Changes in leaf transmittance correspond to averaged chloroplast positions within the mesophyll and are used to assess chloroplast movements quantitatively. The development of photometric devices for measuring blue-light-induced changes in leaf transmittance offered a convenient approach, however, limited in its throughput and restricted mainly to laboratory use (Shibata, 1958; Walczak & Gabrys, 1980; Yorinao & Kazuo, 1974). Leaf reflectance has been also used to monitor chloroplast movements (Park et al., 1996), (Gorton et al., 1999). A method using measurements of red light reflected from the leaf surface was developed to monitor movements during the growth of *Arabidopsis thaliana* in a growth chamber (Dutta et al., 2015). Analysis of leaf optical properties for several species indicated an important role of the cell shape in the ability of chloroplast to move and subsequently alter the leaf absorptance and scattering (Davis et al., 2011). A detailed study of leaf reflectance has also been performed using *Nicotiana tabacum.* Chloroplast positioning resulted in the highest changes in the leaf reflectance spectrum in the region between 500 and 650 nm. The changes in leaf reflectance were about 2% in the blue and red regions and about 5% in the green region of the light spectrum (Baránková et al., 2016). The works of (Davis et al., 2011), (Baránková et al., 2016) used an integrating sphere to measure changes in diffuse reflectance. While this approach yields invaluable data on the effects of chloroplast movements on the optical properties of the leaves, it shares the limitations of the transmittance-based method in its low throughput and requirement of direct contact with the leaf.

The aims of our work were twofold - to establish a system for high throughput chloroplast relocation monitoring using spectra of white light reflected from leaves and to examine whether changes in chloroplast positioning can confound the assessment of leaf traits from commonly used vegetation indices. We have used multiple mutant lines that exhibit disrupted chloroplast movements to distinguish between changes in leaf spectra that can be ascribed to chloroplast relocations from changes due to other light-induced processes. To acquire reflectance spectra and examine their spatial variability within the leaf, we have used hyperspectral imaging, which is a high-throughput, contactless method.

Vegetation indices are a convenient approach to reduce the possibly redundant information in spectra, focusing on the most relevant wavelengths or bands. Hyperspectral or narrow-band vegetation indices are single values calculated using a combination of wavelengths, which are empirically shown to correlate with the traits of interest. Vegetation indices were developed to examine plant or canopy structure (e.g. fractional cover, green biomass, leaf area index, senesced biomass), as well as biochemical (e.g. the content of water, nitrogen, lignin, cellulose, and pigments: chlorophyll, carotenoids, anthocyanins) and physiological traits (e.g. stress, changes in xanthophyll cycle pigments, fluorescence, leaf moisture) (Thenkabail et al., 2018). In particular, a large group of vegetation indices based on the visible or NIR reflectance (formulae in Supplementary Table S1) is used to assess the chlorophyll content, consisting of the classical NDVI (Rouse et al., 1974), but also SR (Rouse et al., 1974), EVI (Huete et al., 1997), ARVI (Kaufman & Tanre, 1995), RENDVI (Gitelson & Merzlyak, 1994), mRESR (Sims & Gamon, 2002), and a sum green index, (Behmann et al., 2014). VOG1, 2, and 3 link the chlorophyll and water contents (Vogelmann et al., 1993). PRI (Gamon et al., 1992), SIPI (Penuelas et al., 1995), CAR1, and CAR2 (Gitelson et al., 2002) represent carotenoid proportions, while RGRI (Gamon & Surfus, 1999), the anthocyanin-chlorophyll ratio. mRENDEVI (Sims & Gamon, 2002) and PSRI (Merzlyak et al., 1999) are used for stress and senescence estimation. The equations specifying vegetation indices are based on the visible and NIR infrared range of the solar spectrum. In the visible and near-infrared range, several bands were found particularly useful for remote sensing, including 660 – 690 nm (red-absorption maxima), 900 – 925 nm (near-infrared reflection peak), 700 – 720 nm (a portion of red-edge), and 540 – 555 nm (green reflectance maxima), 490 nm centered band (rapid change in slope of the spectra), 520 nm and 575 nm bands (the most rapid positive or negative change in reflectance), 845 nm centered band (the NIR shoulder), and 975 nm centered band (biomass/moisture sensitive) (Thenkabail et al., 2002). The estimates of plant traits based on vegetation indices may be biased due to the effects of other, nonspecific physiological responses on the leaf reflectance, such as senescence (Idso et al., 1980); as well as changes in the chlorophyll and carotenoid content (Tucker, 1977). The values of vegetation indices may be also altered by topography (Chen et al., 2020) or snow cover (Wang et al., 2023). In this work, we have investigated the impact of chloroplast positioning on commonly used vegetation indices and discussed the magnitude of their changes caused by chloroplast movements compared to the scale of changes due to the variation of other traits, reported in the literature. We have analyzed the accuracy of vegetation indices for the detection of chloroplast positioning and identified an index of the normalized difference type, tentatively referred to as a Chloroplast Movement Index, that correlates well with the chloroplast arrangement.

To examine whether a more accurate classification of leaves according to chloroplast arrangement can be obtained using whole spectra of reflectance in the visible range, we have examined the performance of three supervised machine learning procedures in the task of classifying leaves according to the chloroplast positioning based on the reflectance spectra. We employed the support vector machine and two types of neural feed-forward network architectures, the multilayer perceptron and convolutional networks. A mathematically rigorous treatment of both of these types of neural networks can be found in (Caterini & Chang, 2018) and (Berner et al., 2022). The Multilayer Perceptron (Pal & Mitra, 1992) is a classical architecture, consisting of a variable number of densely connected layers of neurons. It has been used already in the early works applying machine learning for the analysis of reflectance spectra and remotely sensed data, as reviewed in (Mas & Flores, 2008). In particular, it has been successfully applied to the detection of fungal infections from leaf spectra (Moshou et al., 2004) and to estimate carotenoid content through inversion of the PROSPECT model (Hao et al., 2023). In convolutional neural networks, densely connected layers resembling the multilayer perceptron are preceded by additional layers that convolve their input with an arbitrary number of kernels. The benefits of convolutional networks include their good generalization (Alzubaidi et al., 2021) so that the network performance is usually not substantially degraded when applied to an unseen dataset. They have recently become a popular tool for analysis of spectral data. Their applications include the prediction of the concentration of photosynthetic pigment through the inversion of leaf optical models (Annala et al., 2020), (Shi et al., 2022) as well as the estimation of nitrogen (Pullanagari et al., 2021) and water content (Zhang et al., 2023) from the leaf reflectance spectra. However, an architecture based on the multilayer perceptron was found to be superior to a convolutional network for the prediction of photosynthetic performance in Japanese beaches based on leaf reflectance (Song & Wang, 2021).

## Materials and methods

### Plants and Growth Conditions

Seeds of *Nicotiana benthamiana, Arabidopsis thaliana* wild-type Col-0, *phot1*: SALK_088841 (Lehmann et al., 2011), the *phot2* (*npl1-1*) mutant (Jarillo et al., 2001), *phot1phot2* (SALK_088841 crossed with *npl1-1*), (Hermanowicz et al., 2019), *jac1-3:* WiscDsLox457-460P9 (Hermanowicz et al., 2019) were sown in Jiffy-7 pots (Jiffy Products International AS) and vernalized at 4°C for two days. All plants were transferred to a growth chamber (Sanyo MLR 350H) at 23°C, 80% relative humidity, with a photoperiod of 10 h light and 14 h darkness. The irradiance at the plane of leaf rosettes was 70 μmol·m^-2^·s^-1^. Light was supplied by fluorescent lamps (FL40SS.W/37, Sanyo, Japan). After 4 weeks *N. benthamiana* were transferred to pots containing soil and moved to a walk-in growth chamber with the photoperiod of 10 h light and 14 h darkness, equipped with white LEDs supplying irradiance of ca. 100 μmol·m^-2^·s^-1^.4-week-old *Arabidopsis* and 6-week-old *N. benthamiana* plants were used for the experiment. Plants were dark-adapted for at least 16 h for all kinds of measurements of chloroplast movements. For microscopic observations, detached leaves were either kept in darkness or irradiated with blue light of 1.6 µmol·m^-2^·s^-1^ or 120 µmol·m^-2^·s^-1^ for 1 h (EPILED, 1 W, 460 nm). For hyperspectral imaging half of the leaf was covered with aluminum foil and the other half was irradiated with the same light irradiance as for confocal microscopy.

### Microscopic observations

Microscopic observations were performed with the Axio Observer.Z1 inverted microscope (Carl Zeiss, Jena, Germany) and the LSM 880 confocal module. The long-distance LCI Apochromat 25x, NA 0.8, objective was used with glycerol immersion. Chloroplast positioning was observed in the palisade parenchyma of wild-type *Nicotiana benthamiana* and *Arabidopsis thaliana* leaves as well as *phot1, phot2, phot1phot2,* and *jac1* mutants. The 633 nm He-Ne laser was used to excite chlorophyll, and emission in the range of 661 - 721 nm was recorded as the magenta channel. Stacks were collected on the upper surface of the leaves. Projection images were calculated from slices corresponding to the first ca. 90 μm of stacks, starting from the upper surface of the epidermis. This range included the epidermis and upper parts of palisade cells.

### Photometric method

Leaf transmittance changes due to chloroplast movements were assessed with the photometric method (Gabryś et al., 2017), using a custom-built photometric setup. Chloroplast responses were induced with blue (peak at 455 nm, M455L4 LED, Thorlabs) actinic light of 1.6 or 120 µmol m^−2^ s^-1^. The red measuring beam was produced by a 660 nm LED (M660L4, Thorlabs) and modulated at 1033 Hz. The beams were collimated and combined with a dichroic mirror (DMLP550, Thorlabs). They were then directed toward a detached leaf, mounted in front of a port of an integrating sphere (IS200-4, Thorlabs). The signal was detected with a photodiode detector (DET100A2, Thorlabs) mounted at another port of the sphere. Leaf transmittance curves were processed with a custom-written Mathematica (Wolfram Research) script. Irradiance was measured at the sample plane with the LI-190R sensor (LI-COR) and Keithley 6485 picoampere meter.

### Hyperspectral imaging

Immediately after blue irradiation, the aluminum foil was removed from the covered leaf half, and hyperspectral images of whole leaves were collected using a Headwall Nano hyperspectral Camera (Nano-Hyperspec VNIR 400-1000 nm) equipped with a 12 mm objective (Cinegon F/1.4/12-0906, C-Mount, Schneider Kreuznach, 400-1000 nm). The camera was mounted on a rotating platform (Sevenoak SK-ebh01 Pro), 30 cm from the table. A halogen lamp (Hedler), located 58 cm from the table, was used as a white light source delivering irradiance of 280 μmol·m^-2^·s^-1^ in the leaf plane for collecting the reflected light in the range of 400 - 1000 nm. The angle of incidence of light with respect to the leaf normal was ca. 10°. All images were taken on a black velvet background. White, diffusely-reflecting PTFE disks (SM05CP2C, Thorlabs) were used as reflectance standards. The extraction of spectra and calculation of the mean difference in reflectance spectra between irradiated and darkened halves was performed using a custom script written in Mathematica 13.3 (Wolfram Research).

### Machine learning procedures for leaf classification

The performance of three types of machine learning procedures, the support vector machine, the multilayer perceptron, and the convolutional network, in the classification of leaves based on reflectance spectra were examined using Mathematica 13.3 (Wolfram Research). Spectra were normalized, truncated to the 400 – 750 nm range, and divided into three classes, corresponding to dark-adapted, low-light, and high-light irradiated leaves. As a data augmentation procedure, spectra were extracted from square blocks (60 by 60 pixels), randomly selected within the leaf area. Training and validation were performed using spectra recorded on *N. benthamiana* leaves. To assess the generalization capability of the trained classifiers, final testing was performed on spectra recorded from *N. benthamiana*, as well as on spectra from *A. thaliana*. The *N. benthamiana* dataset was first split into the test set (25%) and the set used for building classifiers (75%). The latter one was further split into the training set and the validation (held-out) set, using balanced five-fold cross-validation. Cross-validation was repeated ten times, with different random partitioning of the dataset into the folds. The performance of the trained classifiers on the validation set guided the selection of model hyperparameters. For the support vector machine, five kernel types were examined (linear, polynomial, sigmoid, and radial Gaussian function). Multilayer perceptron architectures with three or fewer densely connected layers were examined, with up to 30 neurons per layer and a decreasing number of neurons in the input-output direction. Each layer was followed by a batch normalization layer and a rectified linear unit. For the convolution networks, we examined several combinations of the number of convolution layers (1 to 4), the width of the convolution and max pooling kernels, the number of feature maps, as well as number of densely connected layers. Successive convolutional lavers had decreasing kernel width and increasing number of feature maps, which is a common approach. Rectified linear units were always used for activation functions and batch normalization layers were inserted after each densely connected layer to regularize the trained network. The final layer of neurons in the multilayer perceptron and convolutional networks was connected to a softmax layer. As the modeled response variable was categorical, with three classes, the training consisted of minimizing the negative log-likelihood of the multinomial distribution, corresponding to the cross-entropy function. The training was carried out in 2000 epochs, using the Adam optimizer, with a batch size of 64.

## Results

### Hyperspectral imaging of chloroplast movements in Nicotiana benthamiana

To assess the accuracy of hyperspectral imaging for detecting chloroplast movements, we chose a plant with leaf blades of a considerable size, *Nicotiana benthamiana,* which exhibited substantial chloroplast movements (Davis et al., 2011). Blue light of 1.6 μmol·m^-2^·s^-1^ induced a decrease in leaf transmittance corresponding to chloroplast accumulation, whereas blue light of 120 μmol·m^-2^·s^-1^ increased leaf transmittance in line with the chloroplast avoidance response (Fig. 1. A, B). The changes in chloroplast positions due to blue light irradiation observed under the microscope (Fig. 1. C), were visible in the hyperspectral images of partly irradiated leaves (Fig. 1. F, G). The leaf part irradiated with blue light of 1.6 μmol·m^-2^·s^-1^ for 1 h was darker than the part kept simultaneously in darkness by covering it with an aluminum foil. The leaf part irradiated with blue light of 120 μmol·m^-2^·s^-1^ for 1 h appeared paler than the part kept under the aluminum foil (Fig. 1. F, G). The mean reflectance of dark-adapted leaves was ca. 11% in visible light (400-700 nm). The average difference of reflectance in this range between leaves with chloroplasts in avoidance and accumulation positions was ca. 2.3% (Fig. 1. D, E). Differential leaf reflectance curves were calculated from the spectra recorded for non-irradiated and blue light-irradiated leaf parts (Fig. S1, Fig. 1. D, E). After irradiation with 1.6 μmol·m^-2^·s^-1^, a decrease in differential leaf reflectance curves induced by blue light was observed in the green-yellow region (500-600 nm). Irradiation with blue light of 120 μmol·m^-2^·s^-1^ resulted in a characteristic ‘saddle’ in the green-yellow region (500-600 nm) of differential reflectance curves. In addition, a trough at ca. 684 nm and a peak at ca. 695 nm were visible. One-hour irradiation with high blue light induces quenching of chlorophyll fluorescence (Pfündel et al., 2018). Thus, we may expect that the maxima of the chlorophyll emission spectrum (measured in intact leaves) corresponded to troughs in the high-minus-low-light differential curves. Accordingly, the trough at ca. 684 nm is near a small peak at ca. 685 nm visible in the fluorescence emission spectrum of several angiosperm species (Chappelle et al., 1985), (Agati, 1998). The peak at 695 nm in the differential reflectance spectra appears to correspond to the inflection point between the overlapping peaks in the chlorophyll emission spectrum. As the main peak of the emission spectrum from intact leaves is located at ca. 740 nm, fluorescence quenching likely contributes to the steep decrease in the differential reflectance spectrum visible between 700 nm and 745 nm.

**Fig. 1.**
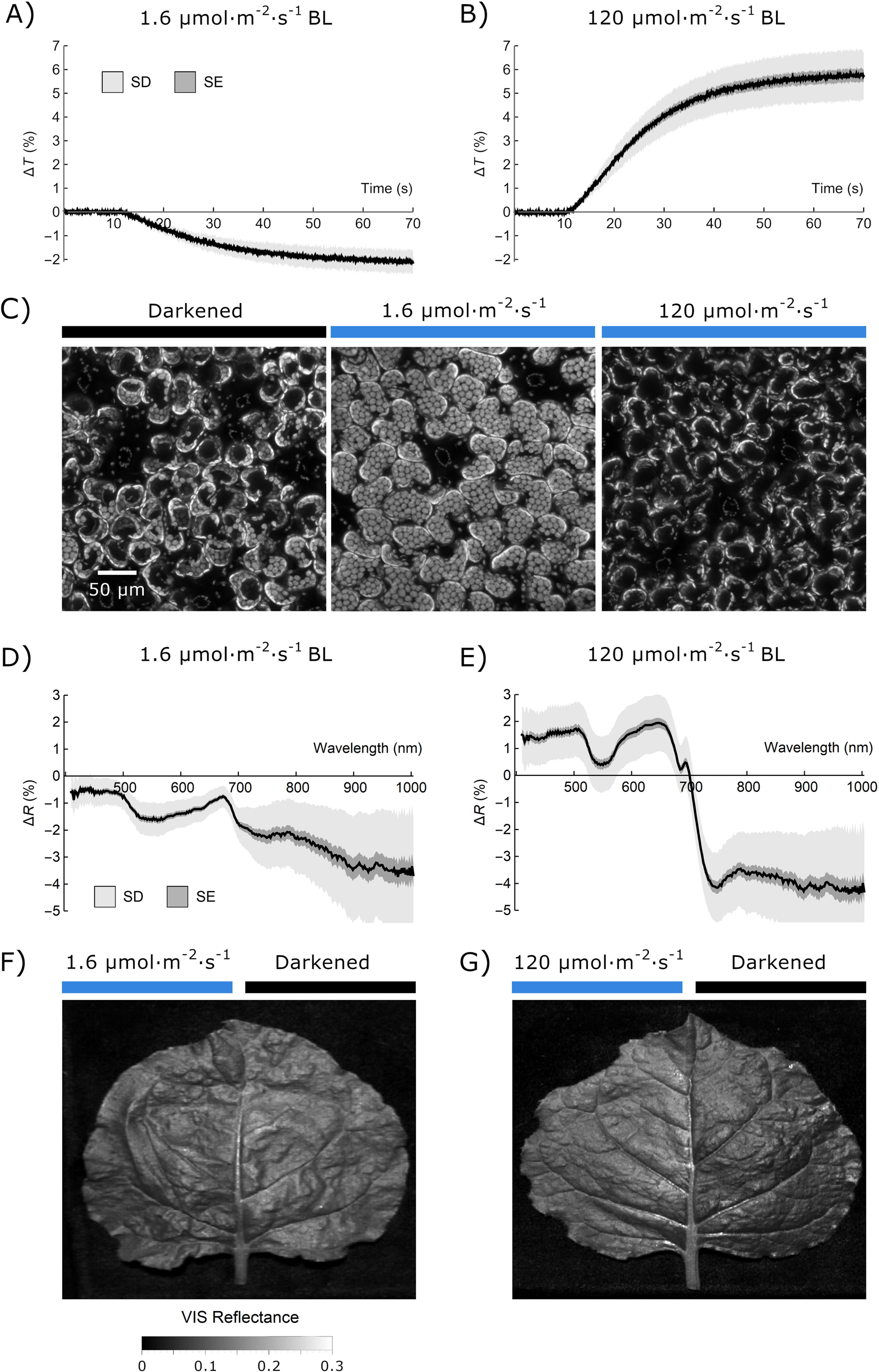
Chloroplast movements in *Nicotiana benthamiana* leaves assessed with (A, B) transmittance measurements, (C) microscopic observations, and (D, E, F, G) leaf reflectance measurements. (A, B) Changes in leaf transmittance at 660 nm, induced by blue light of (A) 1.6 or (B) 120 µmol m^−2^·s^−1^. Each curve represents the mean of 10 measurements. The standard deviation is shown in light grey, the standard error is in dark grey. A decrease in leaf transmittance corresponds to chloroplast accumulation and an increase - to chloroplast avoidance. (C) Chloroplast arrangements in palisade cells of *N. benthamiana*. Leaves detached from 6-week-old dark-adapted plants were either kept in darkness (mock irradiation) or irradiated for 1 h with blue light of 1.6 or 120 µmol·m^−2^·s^−1^. Chloroplast arrangements were imaged with a laser-scanning confocal microscope, using chlorophyll autofluorescence. Maximum intensity projections were calculated from Z-stacks, recorded for ca. 90 μm, starting from the leaf’s upper surface. (D, E, F, G) Effect of blue light irradiation on the reflectance spectrum of *N. benthamiana* leaves. Detached leaves were irradiated with either 1.6 or 120 mmol m^-2^ s^-1^ of continuous blue light (455 nm) for 1 h, with half of the blade covered with aluminum foil. (D, E) The mean difference in reflectance spectra between irradiated and darkened leaf halves calculated from hyperspectral images. The sample size was 23 for low and 31 leaves for high light irradiation. The standard deviation is shown in light grey, the standard error is in dark grey. (F, G) Reflectance images of tobacco leaves, calculated as an average of the hyperspectral images in the visible (400 – 700 nm) range.

### Monitoring chloroplast movements in Arabidopsis thaliana wild type and chloroplast movement mutants

To better understand the relationship between chloroplast positioning and leaf reflectance, we examined light-induced changes in reflectance spectra of several *Arabidopsis thaliana* mutants with disrupted chloroplast movements. The use of mutants was necessary to distinguish between spectral changes due to chloroplast movements and other processes of blue-light-induced responses. Microscopic observations (Fig. 2. A) were performed to enable direct comparisons of the movement aberrations characteristic of mutants. The recorded spectra for *Arabidopsis* wild-type and all chloroplast movement mutants are shown in Fig. S2 & S3. For *Arabidopsis* wild-type plants, patterns of differential leaf reflectance curves were similar to *N*. *benthamiana* (compare Fig 1. D, E and Fig. 2. B). A decrease in the green-yellow region (500-600 nm) after blue light of 1.6 μmol·m^-2^·s^-1^ and a saddle with a peak at ca. 695 nm after blue light of 120 μmol·m^-2^·s^-1^ were observed. Spectra acquired for the *phot1* mutant resembled those of the wild-type, as this mutant differs from the wild-type only at very low light intensities, not investigated in this study. The *phot2* mutant exhibits chloroplast accumulation regardless of the light intensity used, thus the differential leaf reflectance curves were similar for both used light intensities and resembled those of wild-type after blue light treatment of 1.6 μmol·m^-2^·s^-1^. Curves that were representative of the *phot1phot2* mutant did not show specific differences for leaves irradiated with 1.6 μmol·m^-2^·s^-1^ and 120 μmol·m^-2^·s^-1^. The *jac1* mutant, which is characterized by chloroplast avoidance at both investigated light intensities, exhibited curves showing characteristic features for chloroplast avoidance. It is worth noting that the dark positioning in the *jac1* mutant was distinct from the wild-type and *phot1*, which differed from the *phot2* and *phot1phot2* mutants.

**Fig. 2.**
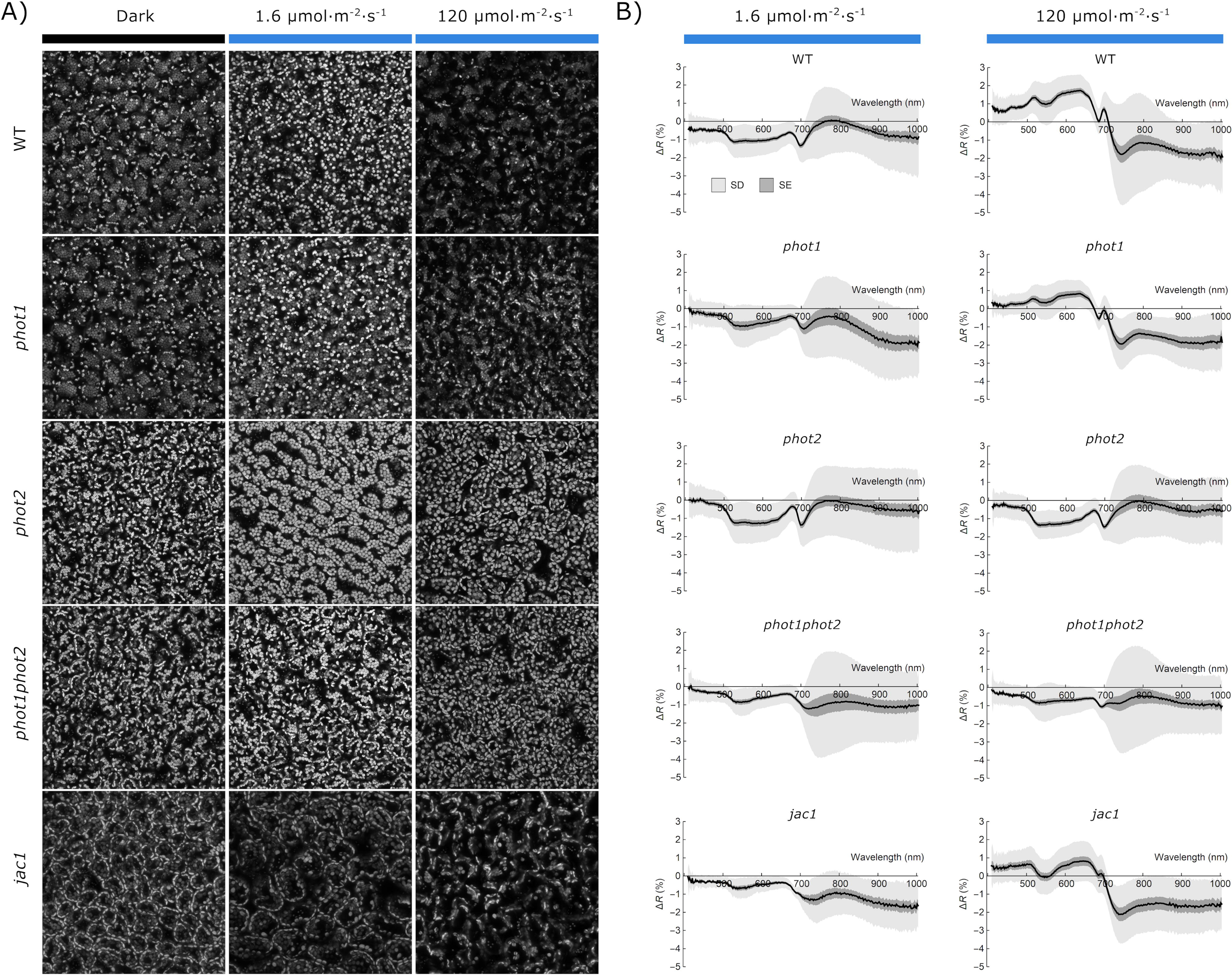
Blue-light-induced chloroplast movements, examined with (A) the optical microscopy and (B) leaf reflectance spectra in *Arabidopsis thaliana* rosette leaves of WT, *phot1*, *phot2*, *phot1phot2,* and *jac1* mutant plants. (A) Chloroplast arrangements in palisade cells of detached leaves from 4-week-old dark-adapted plants were either kept in darkness (mock irradiation) or irradiated for 1 h with blue light of 1.6 or 120 µmol·m^−2^·s^−1^. Chloroplast arrangements were imaged with a laser-scanning confocal microscope, using chlorophyll autofluorescence. Maximum intensity projections were calculated from Z-stacks, recorded for ca. 90 μm, starting from the leaf’s upper surface. (B) Effect of blue light irradiation on the reflectance spectrum of the adaxial surface of *Arabidopsis* wild-type and phototropin mutant leaves. Detached leaves were irradiated with either 1.6 or 120 μmol m^-2^ s^-1^ blue light for 1 h, with half of the blade covered with aluminum foil. The mean difference in reflectance spectra between irradiated and darkened halves was calculated from hyperspectral images. The sample size is 30 leaves for each combination of the plant line and irradiation conditions. The standard deviation is shown in light grey, the standard error is in dark grey.

### Performance of machine learning procedures for classification of leaves according to the position of chloroplast movements

Our next goal was to automatically assess chloroplast positioning using classification algorithms on the spectra extracted from hyperspectral images. Two types of neural networks were considered: a multilayer perceptron and a convolutional neural network (Malek et al., 2018). The multilayer perceptron architecture selected based on the validation set performance consisted of three layers, with 21, 10, and 3 neurons. The structure of the convolutional neural network, selected based on the cross-validation results for multiple trial architectures, is shown in Fig. 3 A. The network consisted of four convolutional layers, interspersed with max-pooling layers and ReLU activation functions. Feature maps generated by the last convolutional layer were concatenated into a single array that acted as an input for densely connected parts of the network, composed of four layers.

**Fig. 3.**
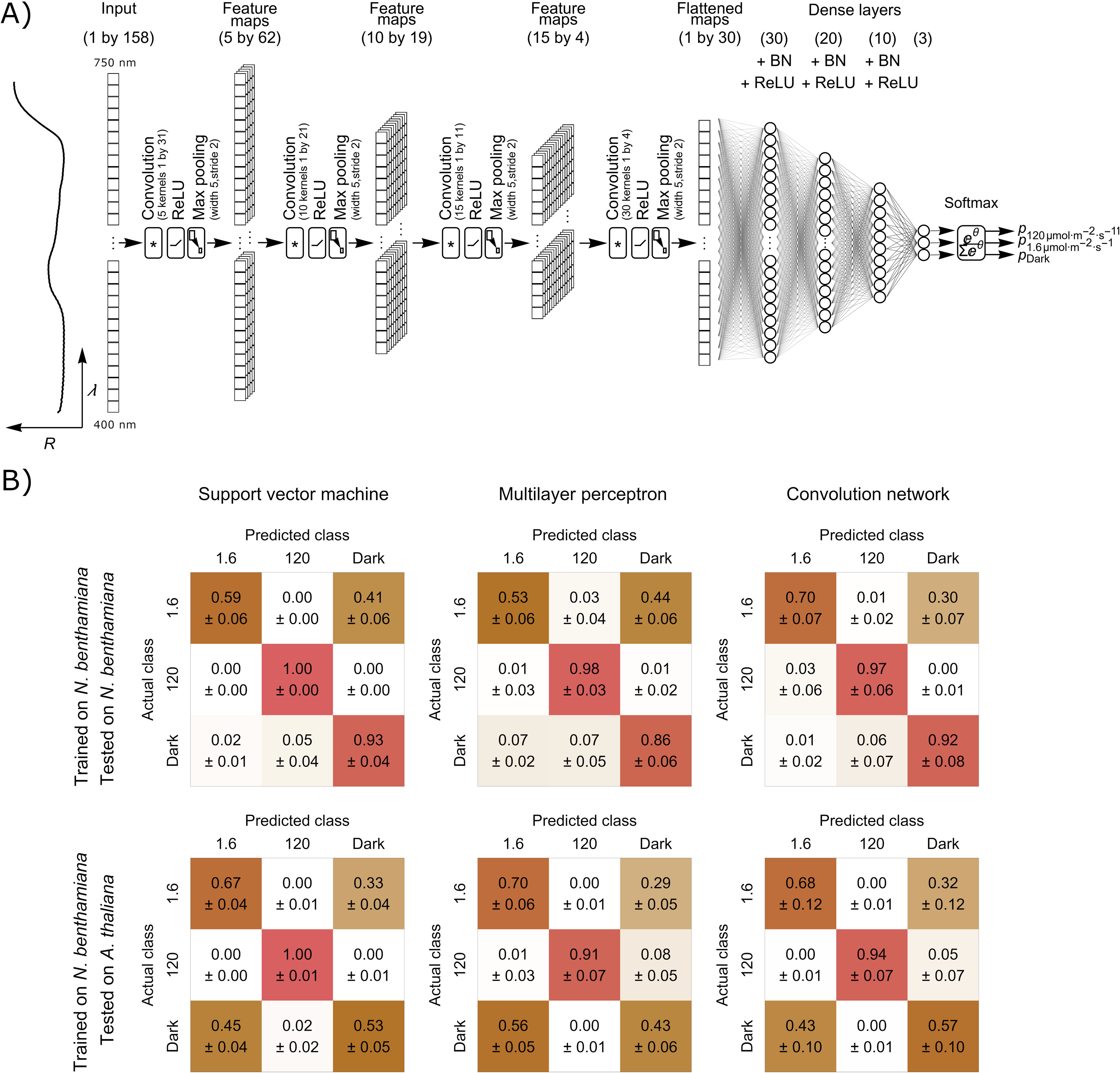
(A) The architecture of the convolutional neuron network for classification of leaf reflectance spectra in the range 400 – 750 nm according to the chloroplast arrangement. The network consists of the initial convolutional part, four densely connected layers, and a softmax layer. The convolutional part consisted of four blocks, each with a 1-D convolution layer of variable kernel size and count, followed by a rectified linear unit (ReLU) and a max pooling layer, with a kernel size of 5 and stride of 2. Each of the first three densely connected layers was followed by a batch normalization (BN) and a ReLU layer. The final softmax layer yields estimated probabilities of the input falling into each chloroplast arrangement class. (B) The confusion matrices showing the mean performance of the support vector machine with a linear kernel, multilayer perceptron, and convolutional network, trained on spectra recorded on *Nicotiana benthamiana* leaves and used to classify spectra recorded for the same species (upper row) or *Arabidopsis thaliana* (bottom row). Training was performed 20 times, and the standard deviation is shown.

All three machine learning methods exhibited very high (91% - 100%) accuracy in distinguishing between spectra recorded on leaves with the chloroplast avoidance positioning and other types of chloroplast positioning (Fig. 3 B). High accuracy was observed both when the trained and test sets came from the same species (*N. benthamiana*) and in the heterologous setup, in which the spectra recorded for *A. thaliana* were used as the test set. The performance of the tested procedures in distinguishing between spectra recorded on leaves exhibiting accumulation and dark positioning was substantially worse than that observed for the avoidance positioning. The best results were obtained using the convolutional network. For the *N. benthamiana* test dataset, 70% of spectra recorded on dark-adapted leaf parts and 92% of spectra recorded on leaves exhibiting the accumulation response were correctly classified.

### Chloroplast Movement Index

To identify wavelengths that can be used to formulate a vegetation index that could be used to distinguish leaves of different chloroplast arrangements, we employed the approach of the correlation matrix, proposed in (Thenkabail et al., 2000). We used the formula of the normalized difference type

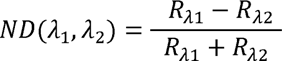

where R_A1_ and R_A2_ are relative reflectance values in narrow bands centered at two wavelengths λ_1_ and λ_2_ in the range of 400 – 750 nm (Fig. 4). For every pair of λ_1_ and λ_2_, the values of the index ND(λ_1_,λ_2_) from raw spectra and its biserial correlation with the chloroplast positioning (binomial variable with two levels) was calculated. The plots were similar for both investigated species, *N. benthamiana* (Fig. 4 A) and *A. thaliana* (Fig. 4 B). We chose the region between 550 nm to 670 nm, in which correlation was high, and identified the local maxima. We additionally took into account that illumination-induced changes in leaf reflectance in the range of 500 nm to 550 nm are likely partly due to changes in chloroplast thylakoid pH gradient and the de-epoxidation of violaxanthin to zeaxanthin (Gamon et al., 1990), thus indices using wavelengths in this spectral region may be biased. Our final formula for the Chloroplast Movement Index, using wavelength marked with a cross in Fig. 4 A and Fig. 4 B, is:

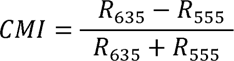

**Fig. 4.**
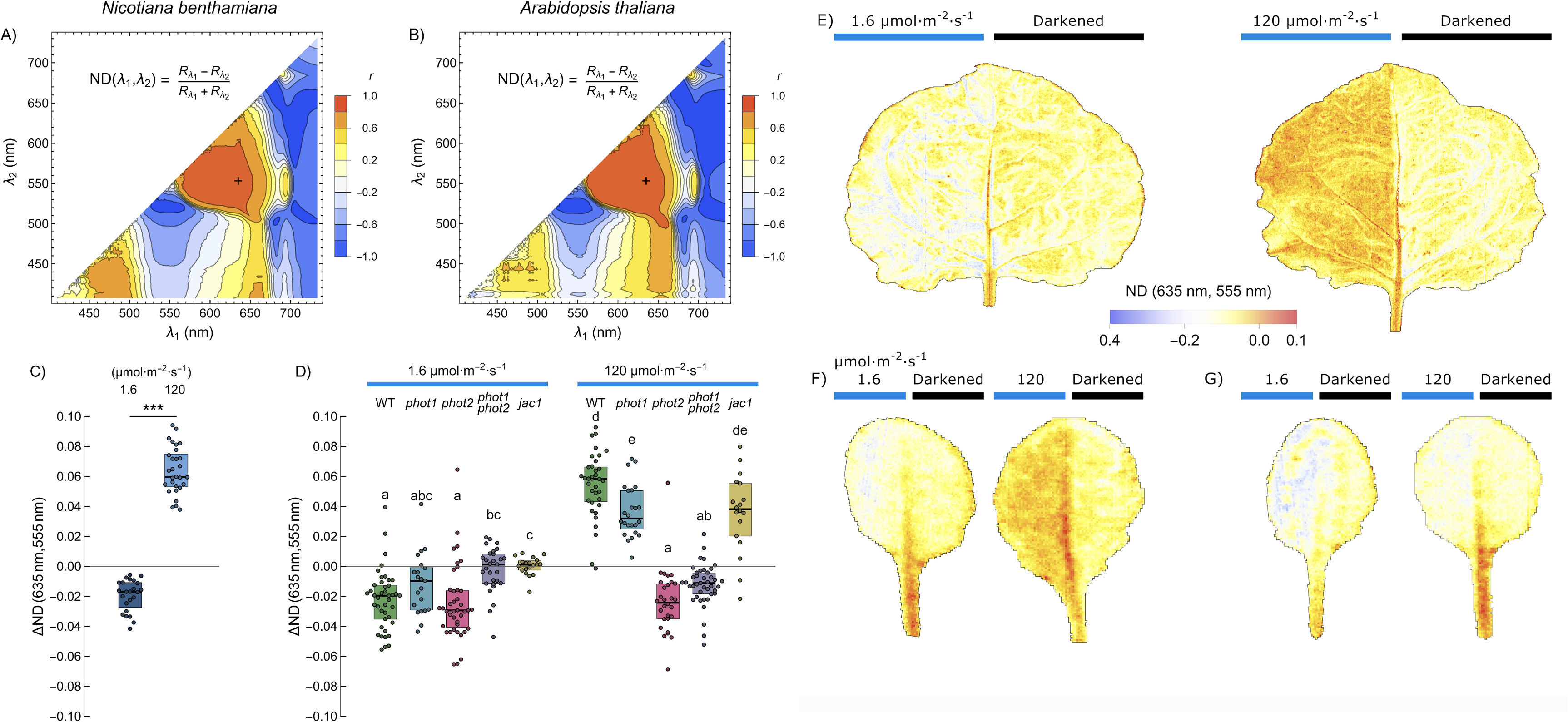
Classification of chloroplast positioning using a vegetation index of a normalized difference type. (A - B) Contour plots showing the biserial correlation between chloroplast positioning and the narrow band vegetation index of the general form specified in the plot, calculated from the leaf reflectance spectra for every combination of wavelengths λ_1_ (x-axis) and λ_2_ (y-axis) in the range of 400 nm to 750 nm. Spectra recorded for (A) *Nicotiana benthamiana* and (B) *Arabidopsis thaliana* leaves were used. Chloroplast accumulation was induced with 1.6 µmol·m^−2^·s^−1^, and avoidance with 120 µmol·m^−2^·s^−1^ of blue light. Blue and red areas correspond to indices that assume high values for leaves exhibiting accumulation and avoidance response, respectively. The pair of wavelengths (635 and 555 nm), used to calculate the ND index for chloroplast movements (CMI), positively correlated with the avoidance response, and was marked with a cross. (C) The difference in the Chloroplast Movement Index between the dark-adapted and either low- or high-light irradiated *N. benthamiana* leaf halves. Each dot corresponds to one leaf. Statistical analysis was performed with the Welch *t*-test. (D) The difference in Chloroplast Movement Index between the dark-adapted and either low- or high-light-irradiated leaf halves of *Arabidopsis thaliana* and chloroplast movement mutants (WT- green*, phot1* - blue*, phot2* – pink, *phot1phot2* - purple*, jac1* - yellow). Boxes that do not share any letter represent groups for which the means of transformed values differ at the 0.05 level (Tukey’s test, adjusted for multiple comparisons). (E, F, G) The Chloroplast Movement Index calculated for each pixel of hyperspectral images of (E) *Nicotiana benthamiana,* (F) *Arabidopsis thaliana* wild-type, and (G) the *phot2* mutant. In each leaf, one half was darkened, while the other was irradiated with either 1.6 µmol·m^−2^·s^−1^ or 120 µmol·m^−2^·s^−1^ of blue light.

We calculated the values of this index for *Nicotiana* (Fig. S4 A) and *Arabidopsis* (Fig. S4 B) spectra and analyzed its performance to distinguish chloroplast accumulation from avoidance. The index took negative values for both species in the range of −0.1 to −0.5. CMI proved to be very effective for *Nicotiana* leaves (Fig. 4 C) irradiated with low and high blue light. For *Arabidopsis* wild type and chloroplast movement mutants (Fig. 4 D), this index seemed to be able to distinguish between chloroplasts in the accumulation and avoidance positions depending on light irradiation regimes, but also the type of mutants. Wild-type plants, *phot1,* and *phot2* mutants in low light and the *phot2* mutant in high light showed chloroplast accumulation. The *phot1phot2* mutant exhibited differences not dependent on chloroplast movements. Wild type, *phot1* and *jac1* mutants in high light demonstrated chloroplast avoidance. We also performed pixel-wise calculations of our Chloroplast Movement Index for whole leaves (Fig.4. E, F, G). Venation and petioles gave high values to the index thus in practice pre-processing of data may be necessary to exclude such areas before analysis, using for example NDVI. In lower-resolution remote sensing spectral unmixing may potentially be used to exclude the contribution of light reflected from tissues other than mesophyll.

### The impact of chloroplast movements on commonly used vegetation indices

We also calculated how chloroplast positioning affects vegetation indices calculated for *Nicotiana benthamiana* leaves. We used a set of indices that are sensitive to pigments and cell structure, as summarized in (Behmann et al., 2014), Supplementary Table S1. Chloroplast positioning after high light treatment reduced the values of several indices, especially NDVI (Fig. 5 A, Fig. S5 A), SR (Fig. 5 B, Fig. S5 B), RENDVI (Fig. 5 F, Fig. S5 F), mRENDEVI (Fig. 5 G, Fig. S5 G), mRESR (Fig. 5 H, Fig. S5 H0, VOG1 (Fig. 5 I, Fig. S5 I), PRI (Fig. 5 L, Fig. S5 L), SIPI (Fig. 5 M, Fig. S5 M), RGRI (Fig. 5 N, Fig. S5 N), PSRI (Fig. 5 O, Fig. S5 O) CAR1 (Fig. 5 P, Fig. S5 P), and CAR2 (Fig. 5 Q, Fig. S5 Q). Chloroplast in the avoidance position elevated SG (Fig. 5 E, Fig. S5 E), VOG2 (Fig. 5 J, Fig. S5 J), and VOG3 (Fig. 5 K, Fig. S5 K) values. Chloroplast accumulation affected vegetation indices to a lesser extent, with diminished SG values and enhanced SR, RENDEVI, mRENDEVI, RGRI, CAR1, and CAR2. EVI (Fig. 5 C, Fig. S5 C) and ARVI (Fig. 5 D, Fig. S5 D) were not dependent on the position of chloroplasts.

**Fig. 5.**
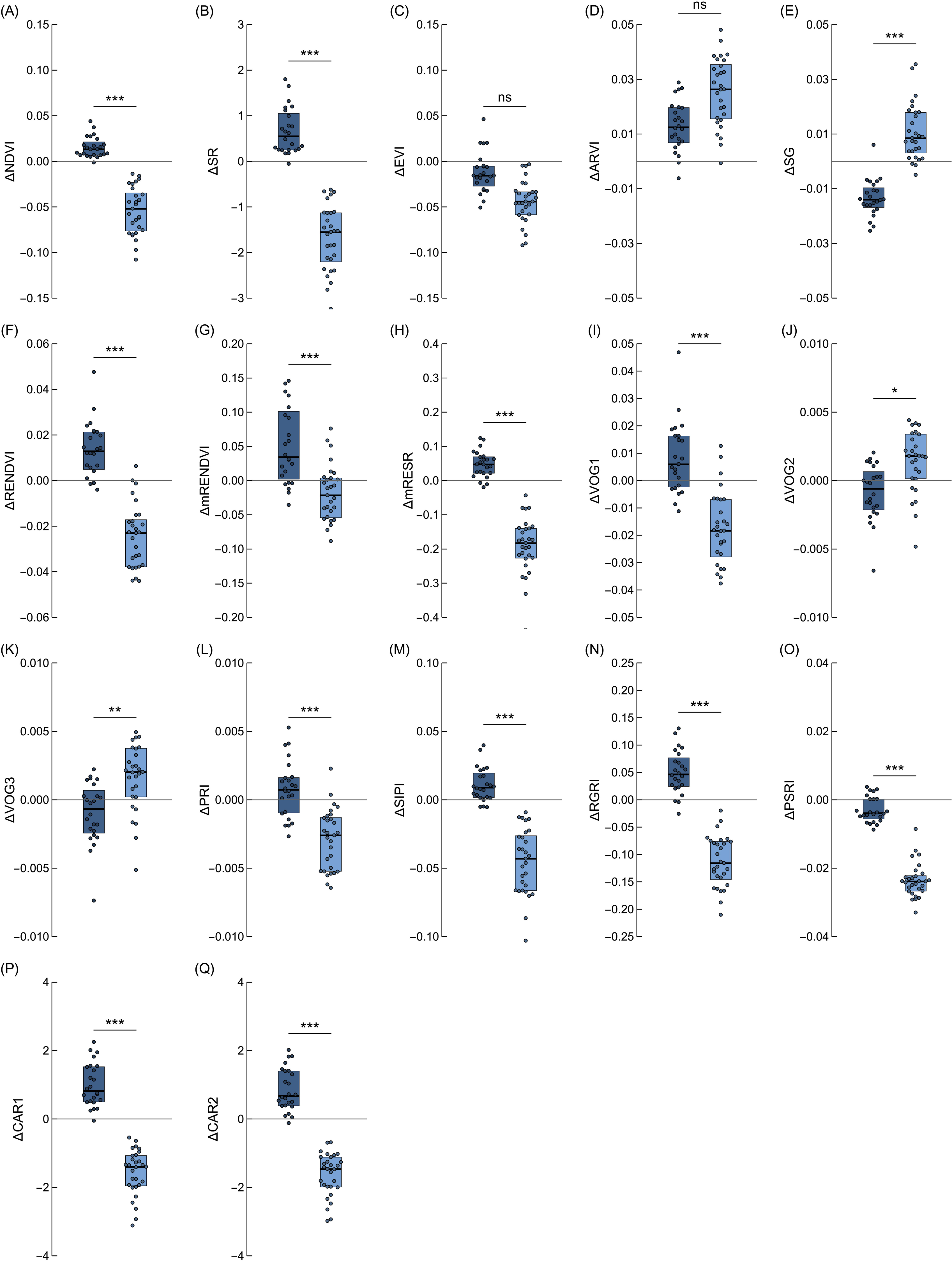
The difference between values of vegetation indices calculated from the spectra from darkened and irradiated parts of the *Nicotiana benthamiana* leaves. Each dot corresponds to one leaf. Irradiation with low blue light of 1.6 µmol·m^−2^·s^−1^ was used to induce chloroplast accumulation (dark blue boxes), with high blue light of 120 µmol·m^−2^·s^−1^ to trigger the avoidance response (light blue boxes). The index formulae are in Supplementary Table 1. The statistical significance of differences between means was examined using the Welch *t-*test.

In wild-type *Arabidopsis,* the direction of change (either elevation or reduction of the index value) was consistent with the results obtained for *Nicotiana* for all investigated indices apart from EVI and PRI (Fig. S5, Fig. S6, Fig. S7, Fig. 5, Fig. 6). The use of *Arabidopsis* chloroplast movement mutants enabled us to determine if the observed changes in the vegetative indices are related to differences in chloroplast arrangements. Values of several indices decreased in leaves showing the avoidance positioning, including NDVI (Fig. 6 A, Fig. S6 A), SR (Fig. 6 B, Fig. S6 B), RENDEVI (Fig. 6 F, Fig. S6 F), mRENDEVI (Fig. 6 G, Fig. S6 G), mRESR (Fig. 6 H, Fig. S6 H), VOG1 (Fig. 6 I, Fig. S7 I), SIPI (Fig. 6 M, Fig. S7 M), RGRI (Fig. 6 N, Fig. S7 N), PSRI (Fig. 6 O, Fig. S7 O), CAR1 (Fig. 6 P, Fig. S7 P) and CAR2 (Fig. 6 Q, Fig. S7 Q). Such a decrease was not observed in mutants deficient in chloroplast avoidance, namely *phot2* and *phot1phot2*. The change in the value of the index mimicked the impact of low light of 1.6 μmol·m^-2^·s^-1^ in those mutants. For indices in which chloroplast avoidance led to an increase in their value, such as SG (Fig. 6 E, Fig. S6 E), VOG2 (Fig. 6 J, Fig. S7 J), and VOG3 (Fig. 6 K, Fig. S7 K), in both mutants, *phot2,* and *phot1phot2* diminished values of those indices were observed, similarly to those observed under low light. RENDEVI, mRENDEVI, and VOG1 seemed to be sensitive to chloroplast accumulation. Changes in EVI (Fig. 6 C, Fig. S6 C), ARVI (Fig. 6 D, Fig. S6 D), and PRI (Fig. 6 L, Fig. S7 L) did not correspond to chloroplast positioning in cells, as no differences were observed between the *Arabidopsis* wild-type and chloroplast movement mutants.

**Fig. 6.**
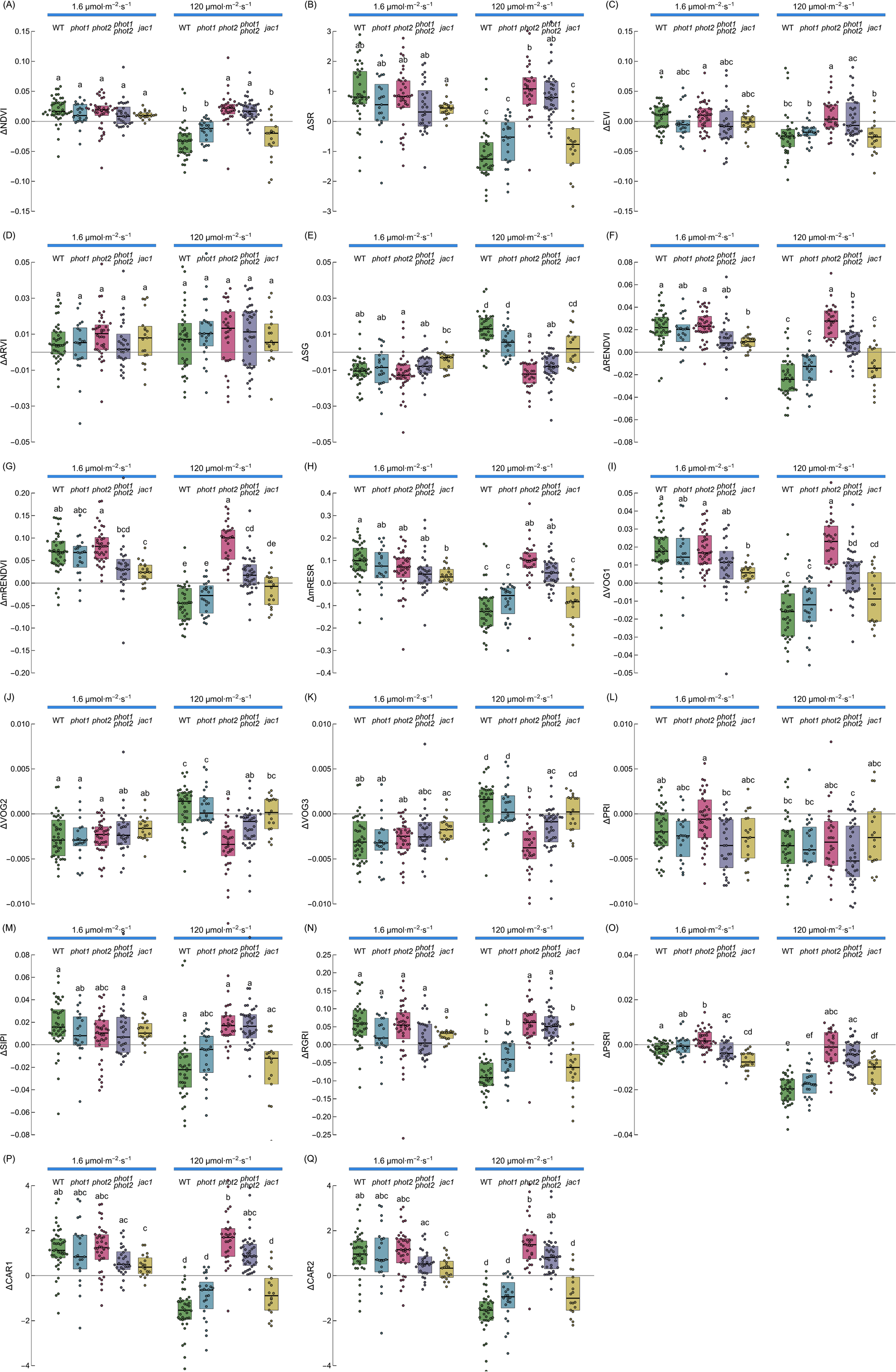
The difference between values of vegetation indices calculated from the spectra from dark-adapted and irradiated parts of *Arabidopsis thaliana* wild-type and chloroplast movement mutants (WT - green*, phot1* - blue*, phot2* - pink, *phot1phot2* - purple*, jac1* - yellow). Each dot corresponds to one leaf. The irradiance to induce chloroplast accumulation was 1.6 µmol·m^−2^·s^−1^ and to trigger chloroplast avoidance was 120 µmol·m^−2^·s^−1^ of blue light. The index formulae are in Supplementary Table 1. Boxes that do not share any letter represent groups for which the means of transformed values differ at the 0.05 level (Tukey’s test, adjusted for multiple comparisons).

## Discussion

### Assessment of chloroplast movements through analysis of hyperspectral images

Here we propose a new method of chloroplast movement assessment based on broad-spectrum reflectance imaging and extraction of information about chloroplast positioning contained in the reflectance spectrum, using either vegetation indices or machine-learning procedures. This method appears to be more suited for high throughput detection of chloroplast movements than methods employing hemispherical reflectance (Davis et al., 2011), (Baránková et al., 2016), which require direct contact of the measuring equipment with the leaf. In the work of (Dutta et al., 2015), directional leaf reflectance of red light was used to visualize chloroplast movements in whole *Arabidopsis* rosettes. The use of white light as the measuring light allowed us to examine the effects of chloroplast positioning not only on the absolute reflectance but also on the shape of the spectrum and ratios between reflectance in selected bands, properties that are often more robust against artifacts or confounding factors (see e.g. (Gamon & Surfus, 1999) in the context of pigment content estimation). These issues are relevant for field conditions when sunlight is used as measuring light.

Averaged hyperspectral images of partly irradiated leaves show a clear difference in the magnitude of visible light reflectance between the irradiated and non-irradiated parts (Fig. 1 F). A work on maize suggested that an increase in PAR reflectance resulting from water stress might stem from the chloroplast avoidance movement (Zygielbaum et al., 2012). Irradiation with blue light affects the shape of the reflectance spectrum (Fig. 1 D, E). The measurements performed on *Arabidopsis* mutants with disrupted chloroplast movements indicate that the observed blue-light-induced changes in the spectrum are to a large extent caused by chloroplast relocations (Fig 2). This suggests that it might be possible to use the information contained in the reflectance spectrum to classify leaves according to the chloroplast positioning. We found that machine learning procedures, in particular the convolution network, exhibited good overall performance in the classification of reflectance spectra of leaves according to the chloroplast positioning (Fig. 3). The support vector machine and neural networks can reliably distinguish leaves exhibiting chloroplast avoidance positioning from the dark-adapted and low-light treated leaves, even when data used for training and testing come from different species (*N. benthamiana* and *A. thaliana*). This indicates that it may be possible to detect chloroplast avoidance even in spectra recorded on heterogeneous vegetation patches.

The Chloroplast Movement Index proposed in this work is the first effort to establish vegetation indices relevant for monitoring chloroplast movements in field conditions (Fig. 4). The differences between values of the index for dark-adapted and irradiated leaves for both investigated species *Arabidopsis thaliana* and *Nicotiana benthamiana* are substantial. The differences of smaller magnitude are observed between dark-adapted and low-light-irradiated leaves. The index characterizes correctly chloroplast positions in *Arabidopsis* mutants with disrupted chloroplast movements and positioning. This suggests that the index is sensitive to chloroplast movements and not to other blue-light-dependent processes. As phototropin mutations affect the leaf morphology and the mesophyll architecture (Kozuka et al., 2011), good performance of the index for the *Arabidopsis* mutants suggests that it may be relatively robust against changes in the leaf structure.

### The impact of chloroplast movements on commonly used vegetation indices

Our results indicate that chloroplast movements may affect several vegetation indices commonly used for remote sensing of physiological traits. To assess the practical significance of this phenomenon, it is necessary to compare the observed changes caused by chloroplast relocations with the magnitude of variation of the indices due to changes in the traits that are usually monitored with them. Analysis of beach forests indicates that vegetation indices are affected by the light conditions in which leaves develop. They generally perform well with sun-lit leaves, but poorly with shaded ones (Sonobe & Wang, 2017). In this work, we show that chlorophyll-dependent indices, characterized by simple formulae, such as Red Edge NDVI (RENDVI), Modified Red Edge NDVI (mRENDVI), Modified Red Edge Simple Ratio Index (mRESR), Simple Ratio Index (SR), as well as Sum Green Index (SG), are affected by chloroplast positioning. The values for RENDEVI or mRENDVI fall in the range of 0 - 0.7 for various leaf types (Sims & Gamon, 2002), and thus may be significantly influenced by the chloroplast relocations. This is especially prominent when differences between dark and lit leaf halves of investigated *Arabidopsis* mutants are analyzed. mRESR values are between 0 - 4 for a variety of leaf types and species. Low values of mRESR, comparable to those calculated in this study, are observed for leaves that are tough or pubescent (e.g. from drought-deciduous species) and are characterized by high surface reflectance (Sims & Gamon, 2002). The Red Green Ratio Index representing the anthocyanin chlorophyll ratio is also substantially affected by chloroplast movements, with ΔRGRI values of about 0.1. RGRI values measured for *Quercus agrifolia* depend on leaf age in the range of 0.5 - 1.5. SR values mainly fall into the range of 0 - 20 as observed for paddy rice fields in Italy and 10 - 20 for clonal *Populus* (Thenkabail et al., 2018). By contrast, the influence of chloroplast positioning is negligible for chlorophyll-dependent indices characterized by more complex formulae, such as the Enhanced Vegetation Index (EVI) and Atmospherically Resistant Vegetation Index (ARVI). The values of EVI, ARVI, SIPI, and PRI seem not to depend on chloroplast positioning, as changes in their values did not correlate in *Arabidopsis* phototropin mutants (Fig. 6). Structure Insensitive Pigment Index (SIPI) values around 1 are shared for a wide variety of plant species, except for evergreen species (Sims & Gamon, 2002). Photochemical Reflectance Index (PRI) values fall in the range of −0.1 to 0.1 except for winter deciduous and evergreen species (Sims & Gamon, 2002). The values of VOG1 and NDVI are very similar between groups both for *Nicotiana* and *Arabidopsis* wild-type and chloroplast movement mutants. The observed differences due to chloroplast positioning are relatively small, although statistically significant. However, NDVI values representing active vegetation are in the range of 0.7 - 0.9 with a calculated uncertainty value of about 0.02 (Borgogno-Mondino et al., 2016). The difference in NDVI between chloroplast dark and avoidance positions is about 0.05, which is significantly above the uncertainty levels for this index.

The accuracy of methods for pigment content estimation from vegetation reflectance spectra is usually lower for carotenoids than chlorophylls (Hill et al., 2019), both at the leaf (Gitelson et al., 2006) and canopy (Asner et al., 2015) levels. This lower accuracy reported for carotenoids is at least partly due to their lower content. However, carotenoid-dependent indices, CAR1 and CAR2, are substantially influenced by chloroplast relocations. For tree species (maple, chestnut, beech) ΔCAR2 of 1 corresponds to a difference in carotenoid content of 1.8 nmol/cm^2^ (0.018 mmol/m^2^). A similar, although nonlinear relationship between carotenoid content and ΔCAR1 is observed for maple leaves (Gitelson et al., 2002). For a survey investigating leaves from a wide range of species representing different functional types, carotenoid content fell in the range of 0 - 0.5 mmol/m^2^ (Sims & Gamon, 2002). Plant Senescence Reflectance Index (PSRI), useful for monitoring stress and senescence, is also significantly influenced by chloroplasts in the avoidance position. The difference that stems from chloroplast positioning may affect PSRI readings, as ΔPSRI values are in the range of 0.02. Values between −0.05 and 0.05 are observed for several annual, deciduous, and evergreen species (Sims & Gamon, 2002).

## Conclusions

We present a contactless method of chloroplast movement detection through hyperspectral imaging. It serves as a proof of concept for a high throughput, remote technique applicable to field conditions. We also propose a new vegetation index of a normalized difference type, useful for assessing the positions of chloroplasts and distinguishing between the accumulation and avoidance responses. Finally, we draw the attention of the community to the impact that chloroplast movement may exert on remote sensing studies based on leaf reflectance. Several commonly used vegetation indices might be affected by the chloroplast positioning in the mesophyll cells.

## Acknowledgements

This research did not receive any specific grant from funding agencies in the public, commercial, or not-for-profit sectors.

## Author contributions

P.H. - Conceptualization; Data curation; Formal analysis; Investigation; Methodology; Software; Validation; Visualization; Writing – original draft, review & editing. J.L. - Conceptualization; Data curation, Validation; Methodology; Investigation; Resources; Supervision; Writing – original draft, review & editing.

## Data availability

The data that supports the findings of this study are available in the supplementary material of this article. Raw data are available from the corresponding author upon reasonable request.

## Supporting Information (brief legends)

Table S1: Formulae of Vegetation Indices calculated from the spectra obtained on *Arabidopsis thaliana* and *Nicotiana benthamiana*

Fig. S1. Leaf reflectance spectra recorded for dark-adapted and illuminated leaf halves of *N. benthamiana.* Detached leaves were irradiated with either (A) 1.6 or (B) 120 mmol m^-2^ s^-1^ of continuous blue light (455 nm) for 1 h, with half of the blade covered with aluminum foil.

Fig. S2. Leaf reflectance spectra recorded for dark-adapted and illuminated leaf halves of *A thaliana* wild-type and chloroplast movement mutants. Detached leaves were irradiated with either 1.6 mmol m^-2^ s^-1^ of continuous blue light (455 nm) for 1 h, with half of the blade covered with aluminum foil.

Fig. S3. Leaf reflectance spectra recorded for dark-adapted and illuminated leaf halves of *A thaliana* wild-type and chloroplast movement mutants. Detached leaves were irradiated with either 120 mmol m^-2^ s^-1^ of continuous blue light (455 nm) for 1 h, with half of the blade covered with aluminum foil.

Fig. S4. Chloroplast Movement Index calculated value for leaf halves of (A) *Nicotiana benthamiana* or (B) *Arabidopsis thaliana* wild type and chloroplast movement mutants (*phot1, phot2, phot1phot2, jac1*) either kept in darkness (grey) or irradiated (blue). The irradiance to induce chloroplast accumulation was 1.6 µmol·m^−2^·s^−1^ and to trigger chloroplast avoidance was 120 µmol·m^−2^·s^−1^.

Fig. S5. Vegetation indices calculated for leaf halves of *Nicotiana benthamiana* either kept in the dark (grey) or irradiated (blue). The formulae for the indices are collected in the Supplementary Table S1. The irradiance to induce chloroplast accumulation was 1.6 µmol·m^−2^·s^−1^ and to trigger chloroplast avoidance was 120 µmol·m^−2^·s^−1^. (A) NDVI, (B) SR, (C) EVI, (D) ARVI, (E) SG, (F) RENDVI, (G) mRENDEVI, (H) mRESR, (I) VOG1, (J) VOG2, (K) VOG3, (L) PRI, (M) SIPI, (N) RGRI, (O) PSRI, (P) CAR1, and (Q) CAR2.

Fig. S6. Vegetation indices calculated for leaf halves of *Arabidopsis thaliana* and chloroplast movement mutants (*phot1, phot2, phot1phot2, jac1*) either kept in the dark (grey) or irradiated (blue). The formulae for the indices are collected in the Supplementary Table S1. The irradiance to induce chloroplast accumulation was 1.6 µmol·m^−2^·s^−1^ and to trigger chloroplast avoidance was 120 µmol·m^−2^·s^−1^. (A) NDVI, (B) SR, (C) EVI, (D) ARVI, (E) SG, (F) RENDVI, (G) mRENDEVI, (H) mRESR.

Fig. S7. Vegetation indices calculated for leaf halves of *Arabidopsis thaliana* and chloroplast movement mutants (*phot1, phot2, phot1phot2, jac1*) either kept in the dark (grey) or irradiated (blue). The formulae for the indices are collected in the Supplementary Table S1. The irradiance to induce chloroplast accumulation was 1.6 µmol·m^−2^·s^−1^ and to trigger chloroplast avoidance was 120 µmol·m^−2^·s^−1^. (I) VOG1, (J) VOG2, (K) VOG3, (L) PRI, (M) SIPI, (N) RGRI, (O) PSRI, (P) CAR1, and (Q) CAR2.

## Abbreviations

ARVI: Atmospherically Resistant Vegetation Index
CAR1: Carotenoid Reflectance Index 1
CAR2: Carotenoid Reflectance Index 2
EVI: Enhanced Vegetation Index
NDVI: Normalized Difference Vegetation Index
mRENDVI: Modified Red Edge NDVI
mRESR: Modified Red Edge Simple Ratio Index
PAR: photosynthetically active radiation
PRI: Photochemical Reflectance Index
PSRI: Plant Senescence Reflectance Index
RENDVI: Red Edge NDVI
RGRI: Red Green Ratio Index
SG: Sum Green Index
SIPI: Structure Insensitive Pigment Index
SR: Simple Ratio Index
VOG1: Vogelmann Red Edge Index 1
VOG2: Vogelmann Red Edge Index 2
VOG3: Vogelmann Red Edge Index 3

