## Supplementary file for "Hyperspectral imaging for chloroplast movement detection"

**Supplementary Data**

Table S1: Formulae of Vegetation Indices calculated from the spectra obtained on *Arabidopsis thaliana* and *Nicotiana benthamiana*

Normalized Difference Vegetation Index (NDVI) – Chlorophyll

$$\frac{R_{800}-R_{670}}{R_{800}+R_{670}}$$

Simple Ratio Index (SR) – Chlorophyll

$$\frac{R_{800}}{R_{670}}$$

Enhanced vegetation Index (EVI) – Chlorophyll

$$2.5\frac{R_{800}-R_{670}}{R_{800}+6R_{670}-7.5R_{490}+1}$$

Atmospherically Resistant Vegetation Index (ARVI) – Chlorophyll

$$\frac{R_{800}-2(R_{670}-R_{490})}{R_{800}+2(R_{670}-R_{490})}$$

Sum Green Index (SG) – Greenness

$$\frac{1}{n}\sum_{i=500}^{599} R_{i}$$

Red Edge NDVI (RENDVI) – Chlorophyll

$$\frac{R_{750}-R_{705}}{R_{750}+R_{705}}$$

Modified Red Edge NDVI (mRENDVI) – Stress, Senescence

$$\frac{R_{750}-R_{705}}{R_{750}+R_{705}-2R_{445}}$$

Modified Red Edge Simple Ratio Index (mRESR) – Chlorophyll

$$\frac{R_{750}-R_{445}}{R_{705}-R_{445}}$$

Vogelmann Red Edge Index 1 (VOG1) – Chlorophyll, Water

$$\frac{R_{740}}{R_{720}}$$

Vogelmann Red Edge Index 2 (VOG2) – Chlorophyll, Water

$$\frac{R_{734}-R_{747}}{R_{715}+R_{726}}$$

Vogelmann Red Edge Index 3 (VOG3) – Chlorophyll, Water

$$\frac{R_{734}-R_{747}}{R_{715}+R_{720}}$$

Photochemical Reflectance Index (PRI) – Photosynthesis, Carot.

$$\frac{R_{531}-R_{570}}{R_{531}+R_{570}}$$

Structure Insensitive Pigment Index (SIPI) – Carot.-Chl.-Ratio

$$\frac{R_{800}-R_{445}}{R_{800}+R_{680}}$$

Red Green Ratio Index (RGRI) – Anth.-Chl.-Ratio

$$\frac{Mean(R_{500-600})}{Mean(R_{600-700})}$$

Plant Senescence Reflectance Index (PSRI) – Stress, Senescence

$$\frac{R_{680}-R_{500}}{R_{750}}$$

Carotenoid Reflectance Index 1 (CAR1) – Carotenoids

$$\frac{1}{R_{510}}-\frac{1}{R_{550}}$$

Carotenoid Reflectance Index 2 (CAR2) – Carotenoids

$$\frac{1}{R_{510}}-\frac{1}{R_{700}}$$

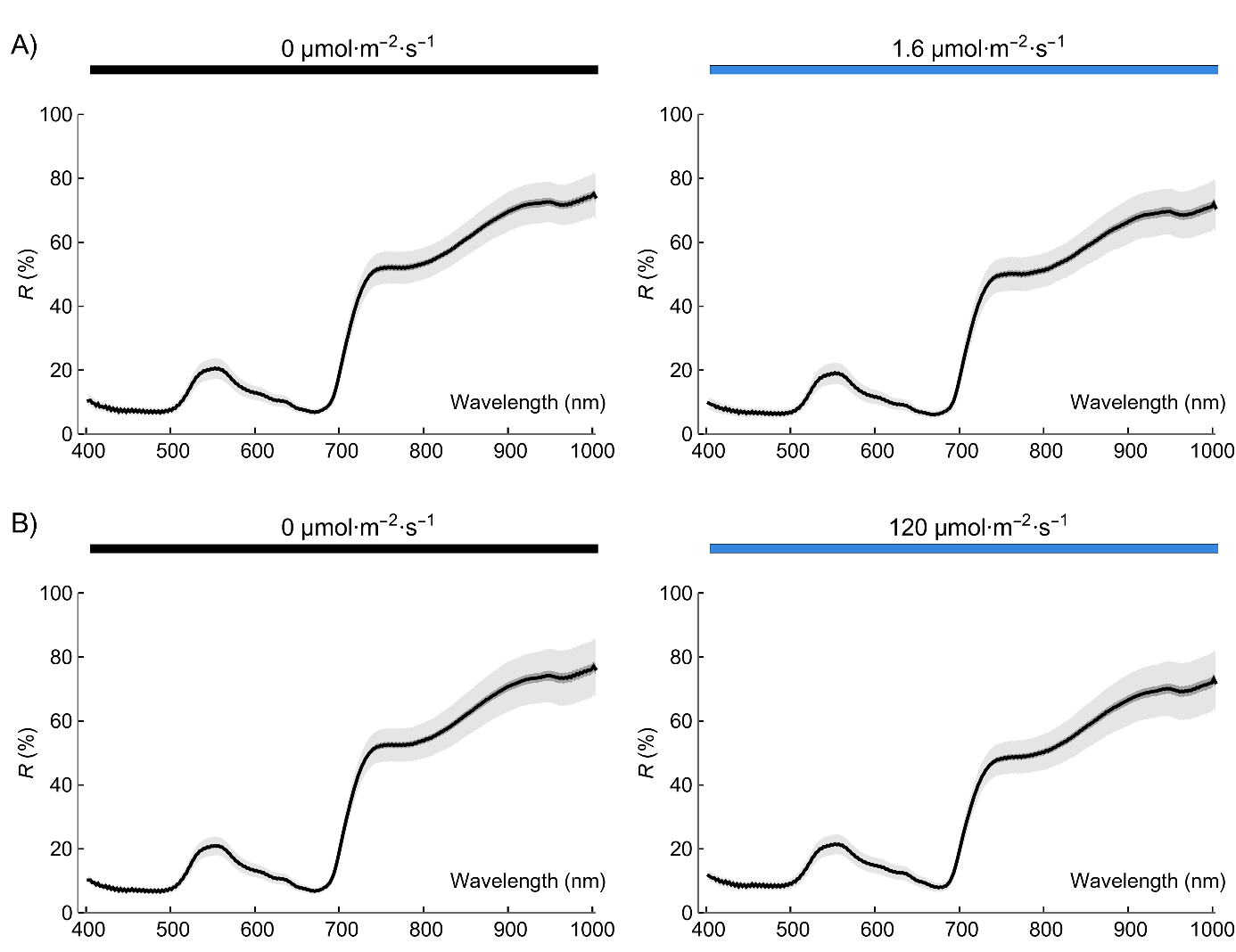

Fig. S1. Leaf reflectance spectra recorded for dark-adapted and illuminated leaf halves of *N. benthamiana.* Detached leaves were irradiated with either (A) 1.6 or (B) 120 mmol m^-2^ s^-1^ of continuous blue light (455 nm) for 1 h, with half of the blade covered with aluminum foil.

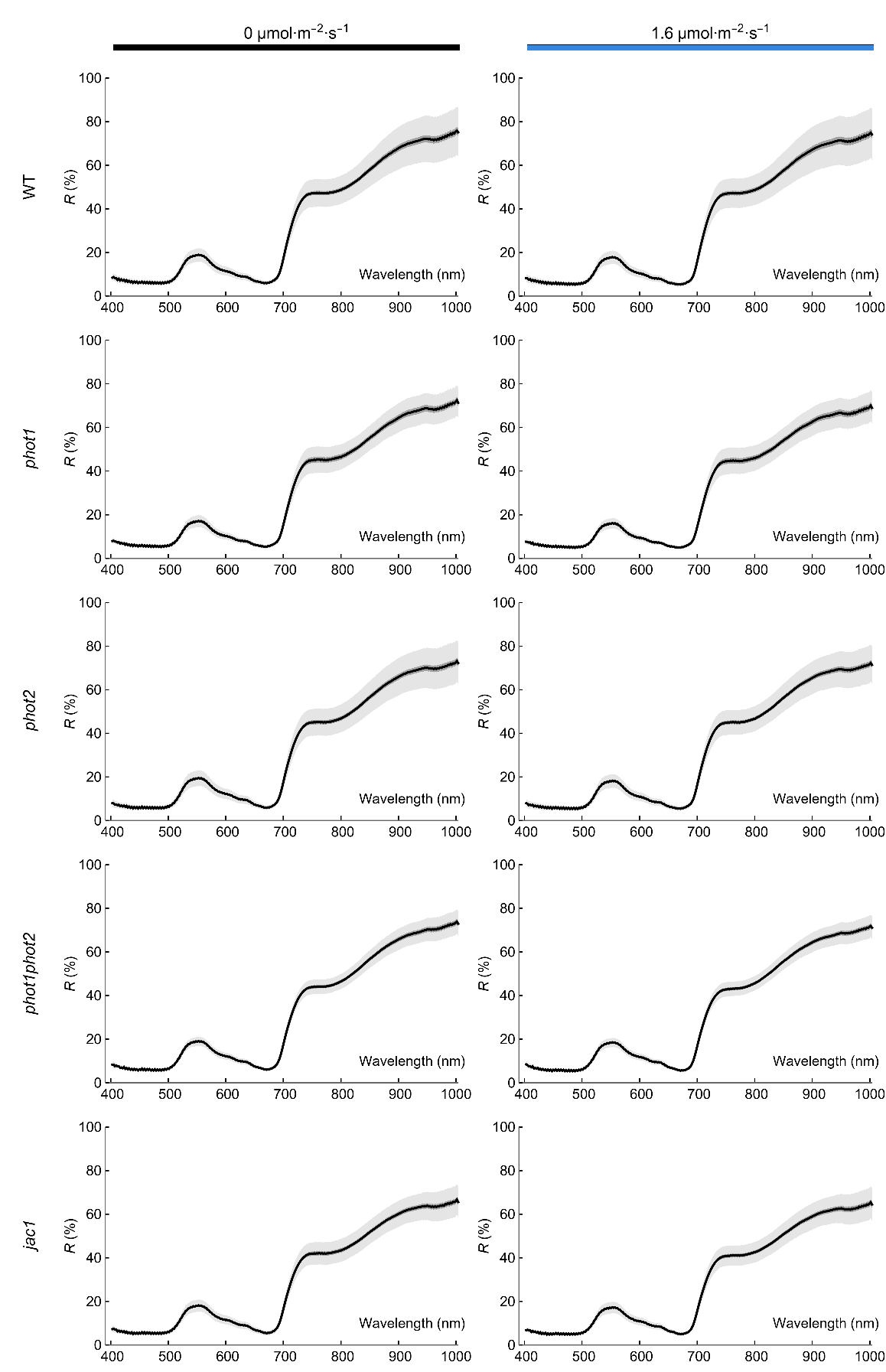

Fig. S2. Leaf reflectance spectra recorded for dark-adapted and illuminated leaf halves of *A thaliana* wild-type and chloroplast movement mutants*.* Detached leaves were irradiated with either 1.6 mmol m^-2^ s^-1^ of continuous blue light (455 nm) for 1 h, with half of the blade covered with aluminum foil.

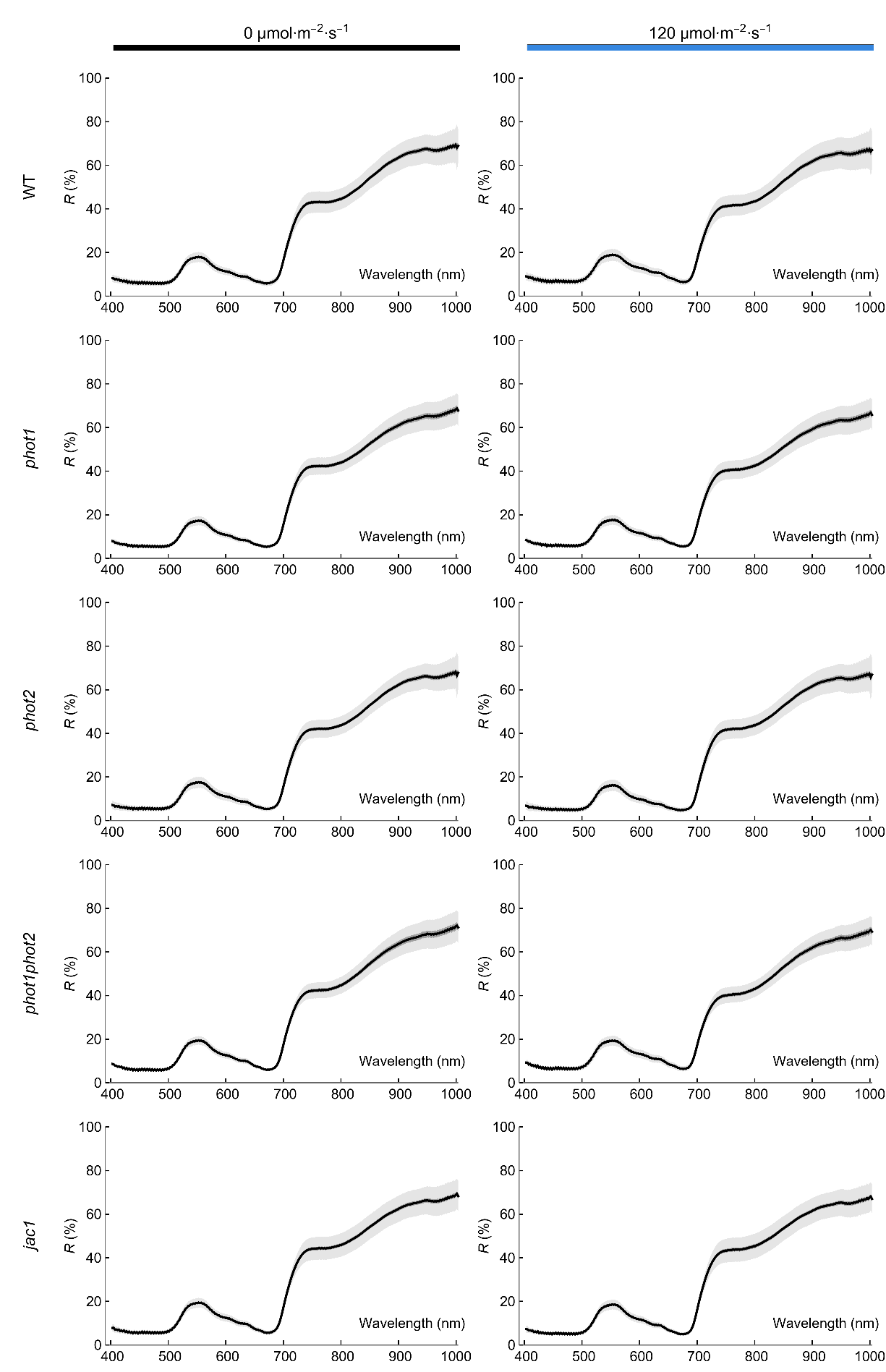

Fig. S3. Leaf reflectance spectra recorded for dark-adapted and illuminated leaf halves of *A thaliana* wild-type and chloroplast movement mutants*.* Detached leaves were irradiated with either 120 mmol m^-2^ s^-1^ of continuous blue light (455 nm) for 1 h, with half of the blade covered with aluminum foil.

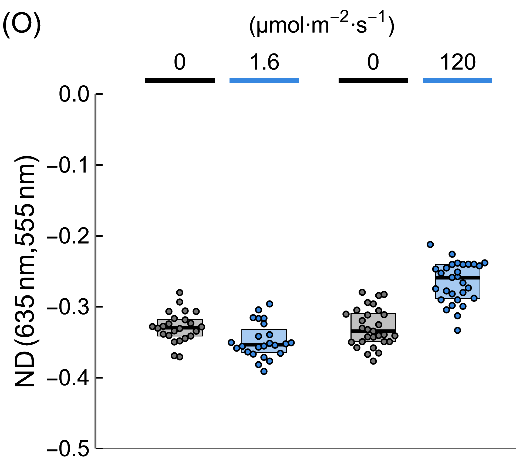

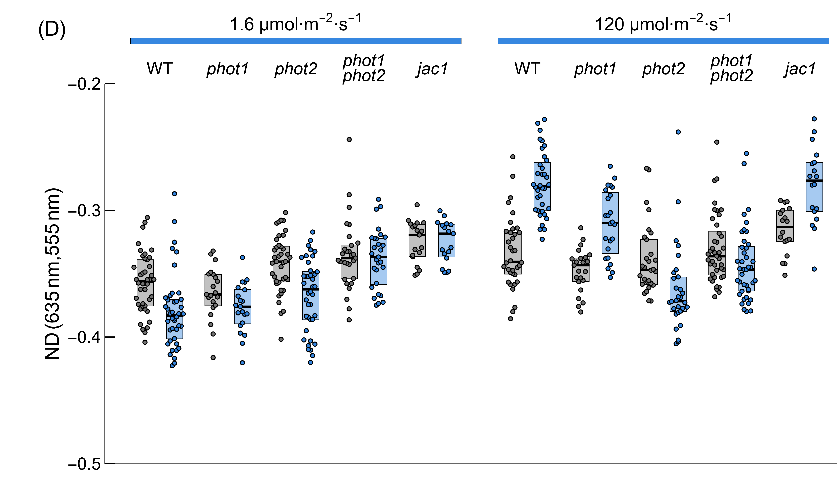

Fig. S4. Chloroplast Movement Index calculated value for leaf halves of (A) *Nicotiana benthamiana* or (B) *Arabidopsis thaliana* wild type and chloroplast movement mutants (*phot1, phot2, phot1phot2, jac1*) either kept in darkness (grey) or irradiated (blue). The irradiance to induce chloroplast accumulation was 1.6 µmol·m^−2^·s^−1^ and to trigger chloroplast avoidance was 120 µmol·m^−2^·s^−1^.

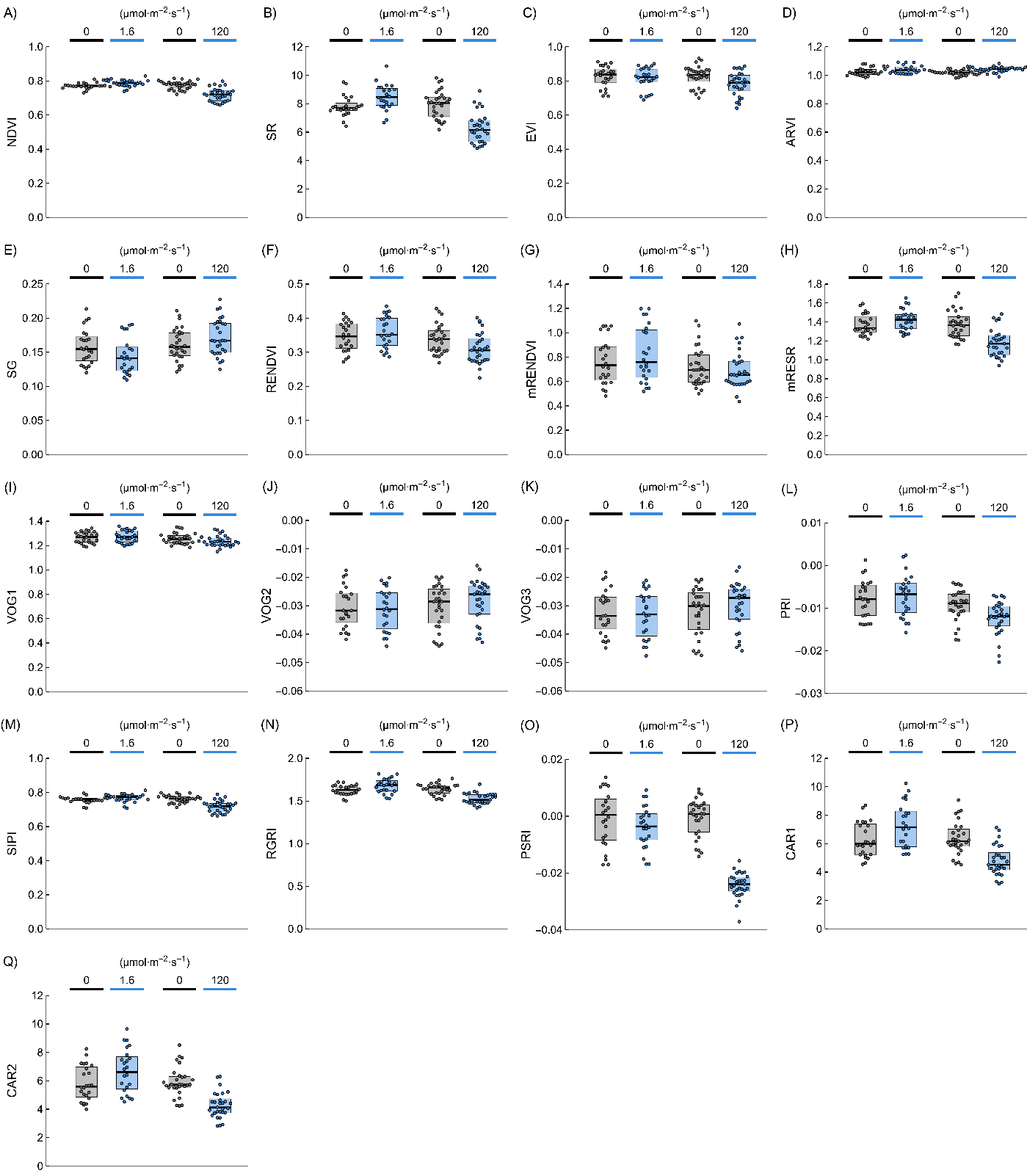

Fig. S5. Vegetation indices calculated for leaf halves of *Nicotiana benthamiana* either kept in the dark (grey) or irradiated (blue). The formulae for the indices are collected in the Supplementary Table S1. The irradiance to induce chloroplast accumulation was 1.6 µmol·m^−2^·s^−1^ and to trigger chloroplast avoidance was 120 µmol·m^−2^·s^−1^. (A) NDVI, (B) SR, (C) EVI, (D) ARVI, (E) SG, (F) RENDVI, (G) mRENDEVI, (H) mRESR, (I) VOG1, (J) VOG2, (K) VOG3, (L) PRI, (M) SIPI, (N) RGRI, (O) PSRI, (P) CAR1, and (Q) CAR2.

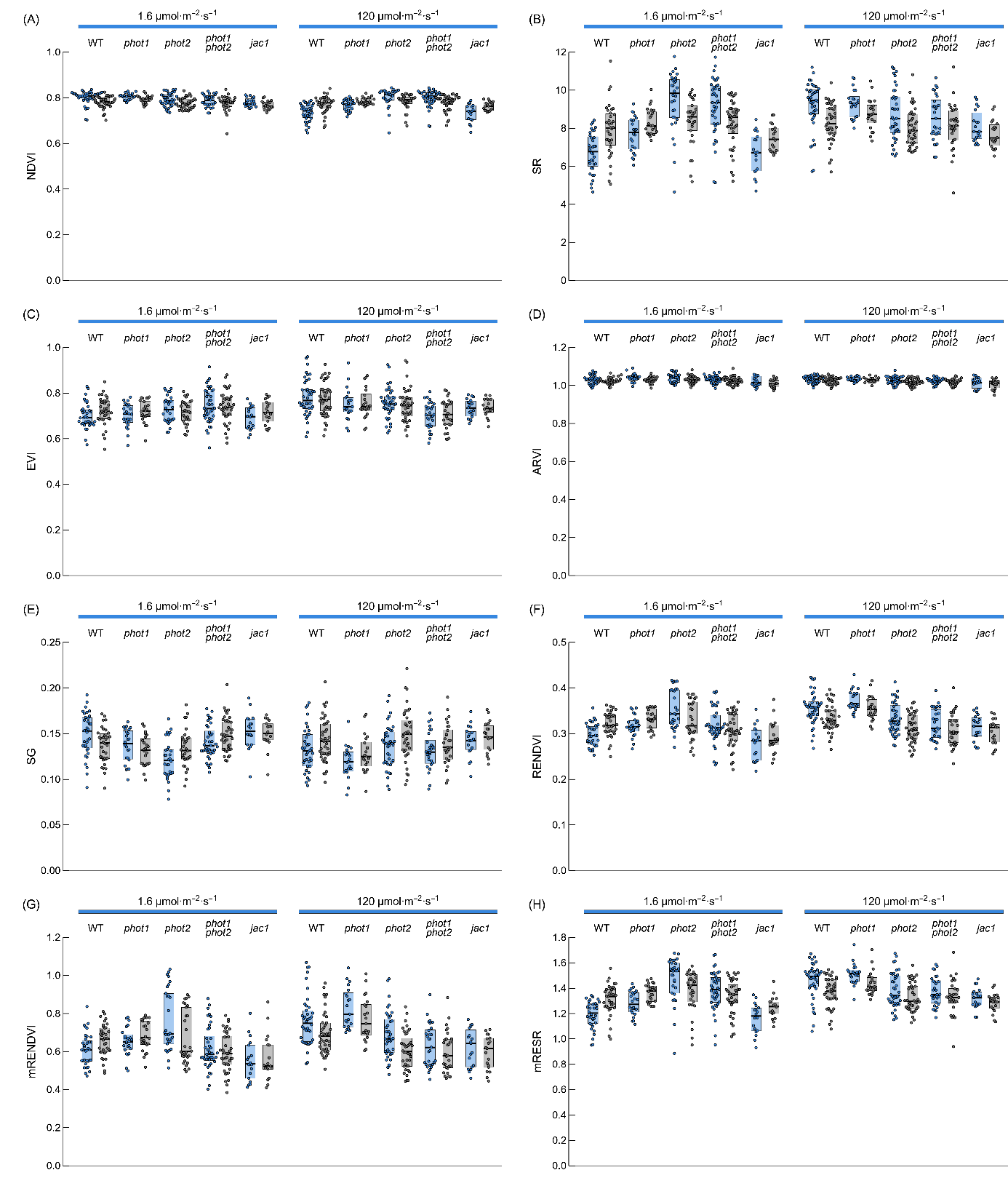

Fig. S6. Vegetation indices calculated for leaf halves of *Arabidopsis thaliana* and chloroplast movement mutants (*phot1, phot2, phot1phot2, jac1*) either kept in the dark (grey) or irradiated (blue). The formulae for the indices are collected in the Supplementary Table S1. The irradiance to induce chloroplast accumulation was 1.6 µmol·m^−2^·s^−1^ and to trigger chloroplast avoidance was 120 µmol·m^−2^·s^−1^. (A) NDVI, (B) SR, (C) EVI, (D) ARVI, (E) SG, (F) RENDVI, (G) mRENDEVI, (H) mRESR.

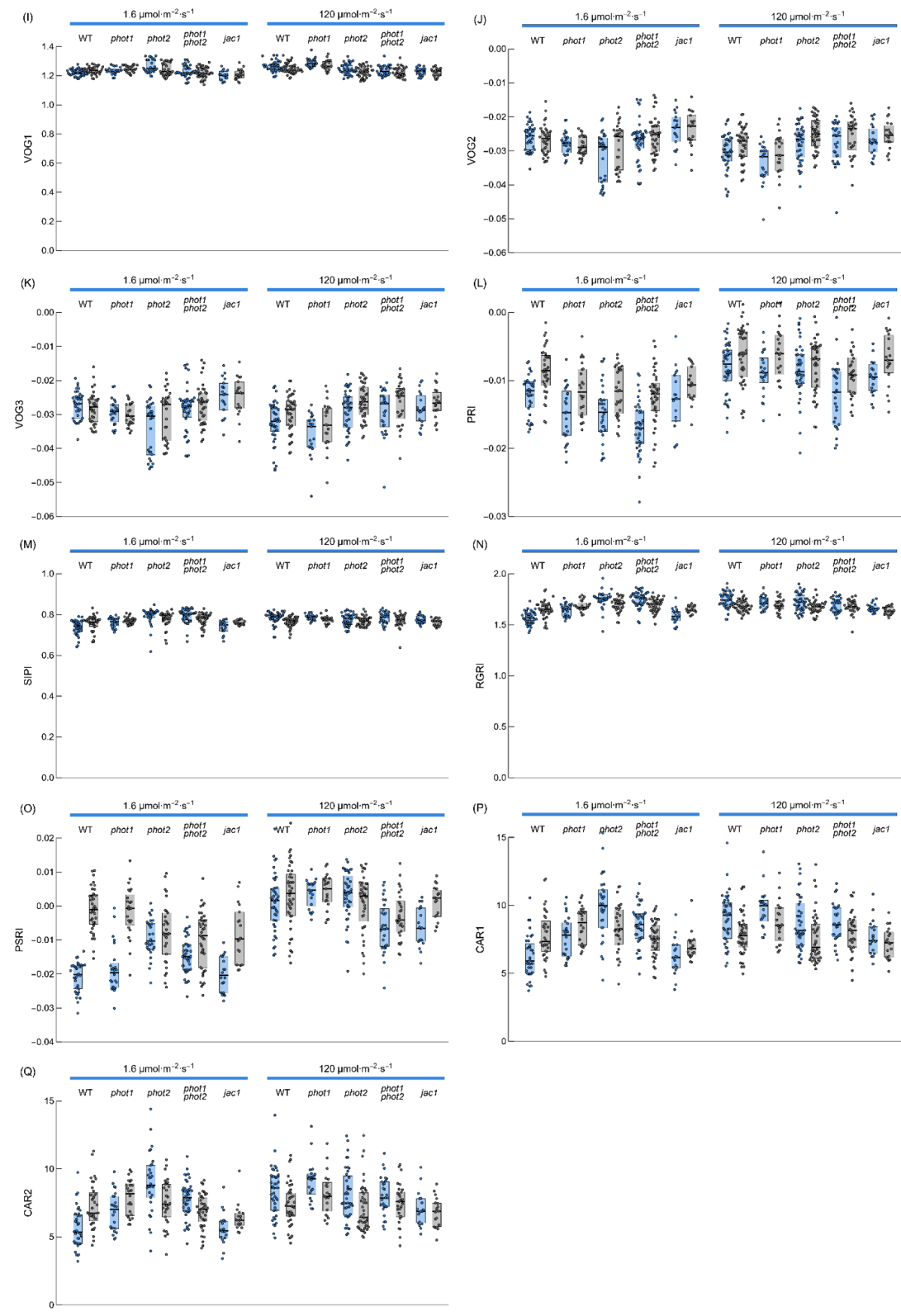

Fig. S7. Vegetation indices calculated for leaf halves of *Arabidopsis thaliana* and chloroplast movement mutants (*phot1, phot2, phot1phot2, jac1*) either kept in the dark (grey) or irradiated (blue). The formulae for the indices are collected in the Supplementary Table S1. The irradiance to induce chloroplast accumulation was 1.6 µmol·m^−2^·s^−1^ and to trigger chloroplast avoidance was 120 µmol·m^−2^·s^−1^. (I) VOG1, (J) VOG2, (K) VOG3, (L) PRI, (M) SIPI, (N) RGRI, (O) PSRI, (P) CAR1, and (Q) CAR2.
